# SARS-COV-2 Omicron variant predicted to exhibit higher affinity to ACE-2 receptor and lower affinity to a large range of neutralizing antibodies, using a rapid computational platform

**DOI:** 10.1101/2021.12.16.472843

**Authors:** Adam Zemla, Thomas Desautels, Edmond Y. Lau, Fangqiang Zhu, Kathryn T. Arrildt, Brent W. Segelke, Shankar Sundaram, Daniel Faissol

## Abstract

Rapid assessment of whether a pandemic pathogen may have increased transmissibility or be capable of evading existing vaccines and therapeutics is critical to mounting an effective public health response. Over the period of seven days, we utilized rapid computational prediction methods to evaluate potential public health implications of the emerging SARS-CoV-2 Omicron variant. Specifically, we modeled the structure of the Omicron variant, examined its interface with human angiotensin converting enzyme 2 (ACE-2) and evaluated the change in binding affinity between Omicron, ACE-2 and publicly known neutralizing antibodies. We also compared the Omicron variant to known Variants of Concern (VoC). Seven of the 15 Omicron mutations occurring in the spike protein receptor binding domain (RBD) occur at the ACE-2 cell receptor interface, and therefore may play a critical role in enhancing binding to ACE-2. Our estimates of Omicron RBD-ACE-2 binding affinities indicate that at least two of RBD mutations, Q493R and N501Y, contribute to enhanced ACE-2 binding, nearly doubling delta-delta-G (ddG) free energies calculated for other VoC’s. Binding affinity estimates also were calculated for 55 known neutralizing SARS-CoV-2 antibodies. Analysis of the results showed that Omicron substantially degrades binding for more than half of these neutralizing SARS-CoV-2 antibodies, and for roughly 10 times as many of the antibodies than the currently dominant Delta variant. This early study lends support to use of rapid computational risk assessments to inform public health decision-making while awaiting detailed experimental characterization and confirmation.

## Background

The recently emerged Omicron variant of SARS-CoV-2 raised significant concerns on whether this variant, compared with the currently dominant Delta variant, will increase transmissibility and degrade immunity from vaccines or protection from neutralizing antibodies. This concern is primarily driven by an exceptionally large number of mutations present on the RBD, including several mutations that have not previously been observed in widely circulating variants. Stabilized expression of this substantially modified spike/RBD followed by in-vitro assessment of affinity and neutralization to antibodies and receptors is the gold standard investigational avenue, but takes several weeks even with the heightened urgency surrounding this variant.

### Objective

On November 26, prior to experimental data/results being made public, we initiated an effort to apply our computational prediction platform to address the following questions.

1. Does Omicron affect affinity to ACE-2, with implications for transmissibility and infectivity?
2. Compared to previous SARS-CoV-2 variants, does Omicron affect/alter recognition by antibodies generated via vaccination or infection/exposure, with implications for adequacy of current countermeasures (monoclonal antibodies and vaccines)?

Rapid availability of reliable predictions can help provide grounding data for decisions on preparedness and tailored response to emerging variants.

## Methods

We have been developing computational methods to rapidly construct structural models of modified RBD’s and predict affinity to ACE-2 or neutralizing antibodies. A manuscript thoroughly describing our method, together with experimental validation demonstrating its accuracy, is in preparation and will be publicly available at a future date. The platform is designed to provide robust and reproducible computational predictions over a large number of binding complexes (multiple variant RBDs, multiple antibodies), with an interpretable confidence estimate, in a matter of days.

The information we utilized to make our predictions includes sequence information for the Omicron variant (no structures are currently available) and the RBD-Fab complexes of publicly available antibodies deposited in the Protein Data Bank (PDB). Our approach, which we call Structural Fluctuation Estimation (SFE), starts with the construction of structural models for each of the RBD-ACE2 and RBD-Fab complexes. For each constructed complex we performed minimization and relaxation using disparate approaches: Rosetta [1], Chimera [2], GROMACS [3] steepest decent and conjugant gradient minimizations, and molecular dynamics trajectories from GROMACS simulations. This procedure allowed us to sample a large number of structural conformations for each RBD-Fab and RBD-ACE2 complex. Using high performance computing systems at the Lawrence Livermore National laboratory for all calculated conformations, we performed “forward” and “reverse” ddG calculations using Flex ddG [4], removed outliers and averaged results from the interquartile only. A distribution of binding free energies (ddG’s) was then computed for each complex providing an affinity estimate together with error and confidence levels.

By computing ddG’s across many conformations of each evaluated complex, we reduced uncertainties in the structural modeling and captured natural conformational fluctuations in the complexes. Each of the final ddG estimates for a given complex is a result from evaluation of 60 conformations requiring about 24 hours on a single 16-core CPU system. In our analysis, we use predicted ddG’s as a proxy for predicted binding affinity both to ACE-2 and neutralizing antibodies.

## Results

### Structural model of Omicron, in-complex with ACE-2, reveals potentially significant changes in the RBD-ACE2 interface region

Fig 1 illustrates our constructed structural model of Omicron in complex with ACE-2. Compared to the wild-type Wuhan strain, Omicron has 30+ mutations in the Spike protein (key protein for virus entry into host cells), which is many more than the number of mutations observed in any other variant to date. Fifteen of those Spike protein mutations are in the receptor binding domain (RBD). Analyzing the RBD, we noted that some residues changed charge, which expanded the number of charged residues by 4. Seven of 15 Spike mutations occur in the interface with ACE-2 cell receptor, shown in yellow in Fig 1. By comparison, the Delta variant (and any other variants of concern) show no more than 15 mutations in the Spike with no more than 3 in the RBD and at most one mutation in the interface with ACE-2.

**Fig. 1.**
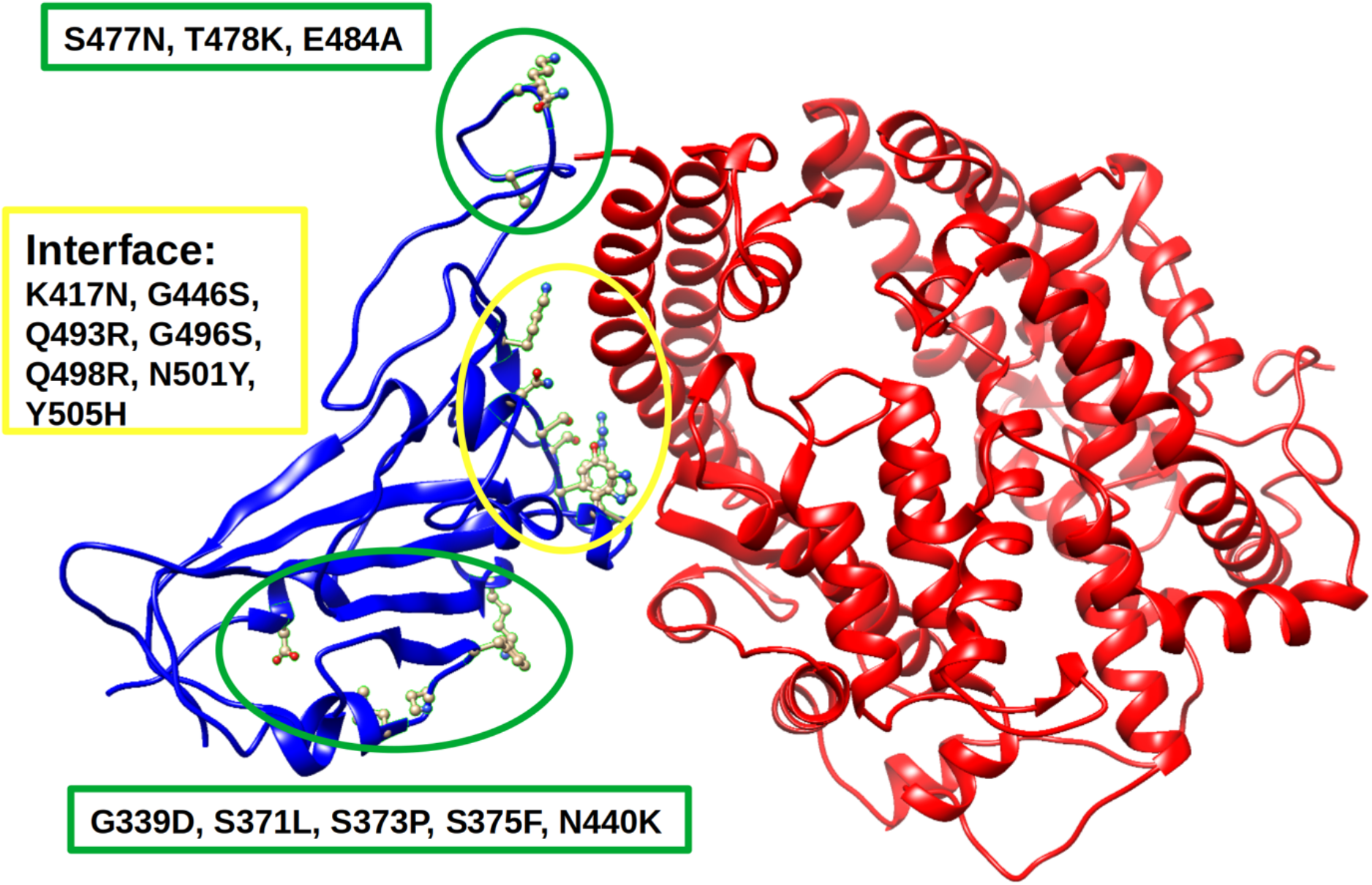
Structural model of SARS-CoV-2 RBD Omicron variant (in blue) in complex with ACE-2 (in red). RBD mutations observed within the 5 Ångstroms interface with ACE-2 are highlighted in yellow, and mutations outside the RBD-ACE-2 interface are highlighted in green.

### Predictions suggest Omicron has increased binding affinity to ACE-2, driven by a few dominant single amino acid changes

To provide insight on whether Omicron has increased affinity to the ACE-2 receptor and thereby potential for increased viral load or transmissibility, we computationally predicted its affinity to ACE-2 relative to Wuhan, Delta and other VoCs. In Fig 2 and Table 1 we provide estimates of binding affinity changes (ddG) using SFE upon introduced mutations to SARS-CoV2 RBD-ACE-2 complexes. SFE predictions indicate that the Omicron variant as well as Omicron with an additional mutation R346K (we call it “Omic346”) have higher affinity (lower binding energy) to ACE-2 than Delta and other VoC’s. Single amino acid change prediction suggests that Q493R and N501Y seem to be the dominant mutations leading to improved ACE-2 binding. In our comparison of Omicron with other VoC’s we added two versions of the Delta variant detected in UK, which we labeled in our tables as “Delta1” (with additional K417N) and “Delta2” (with additional N501Y). Table 1 provides a list of evaluated RBD mutants, including known SARS-CoV-2 VoC’s and some selected single point mutations. Nine of these VoC’s or mutant RBD’s in Table 1, denoted with an “*” were previously experimentally evaluated in [5], and our predictions are consistent with these experimental measurements.

**Table 1.** SARS-COV-2 RBD complexed with the ACE-2 receptor is evaluated for ddG changes upon replacement of the wild-type SARS-CoV-2 RBD with 32 different RBD mutants.

| Variant | SFE (ddG) | Variance | RBD-Mutations |
| --- | --- | --- | --- |
| Alpha | -1.56 | 1.31 | E484K,S494P,N501Y |
| *Beta | -1.23 | 1.85 | K417N,E484K,N501Y |
| Delta | 0.01 | 0.00 | L452R,T478K |
| Delta1 | 0.61 | 0.54 | K417N,L452R,T478K |
| Delta2 | -1.69 | 1.25 | L452R,T478K,N501Y |
| Epsilon | -0.01 | 0.01 | L452R |
| *Eta | 0.07 | 0.02 | E484K |
| *Gamma | -1.13 | 2.01 | K417T,E484K,N501Y |
| Iota | -0.01 | 0.14 | L452R,S477N,E484K |
| Kappa | 0.01 | 0.01 | L452R,E484Q |
| Lambda | -0.03 | 0.03 | L452Q,F490S |
| Mu | -1.47 | 0.84 | R346K,E484K,N501Y |
| Omicron | -2.72 | 1.24 | G339D,S371L,S373P,S375F,K417N,N440K,G446S,S477N,T478K,E484A,Q493R,G496S,Q498R,N501Y,Y505H |
| Omic346 | -2.71 | 1.12 | G339D,R346K,S371L,S373P,S375F,K417N,N440K,G446S,S477N,T478K,E484A,Q493R,G496S,Q498R,N501Y,Y505H |
| *Theta | -1.85 | 1.45 | E484K,N501Y |
| G339D | 0.00 | 0.00 | G339D |
| R346K | 0.00 | 0.00 | R346K |
| *K417N | 0.80 | 0.41 | K417N |
| *K417N,E484K | 0.81 | 0.64 | K417N,E484K |
| *K417T | 0.69 | 0.51 | K417T |
| *S447N | 0.03 | 0.03 | S447N |
| S477N,N501Y | -1.48 | 0.95 | S477N,N501Y |
| Y449H | -0.17 | 0.15 | Y449H |
| Y449H,E484K,N501Y | -1.84 | 1.42 | Y449H,E484K,N501Y |
| T478K | 0.01 | 0.00 | T478K |
| E484A | 0.03 | 0.00 | E484A |
| Q493K | -0.62 | 0.33 | Q493K |
| Q493R | -1.49 | 0.03 | Q493R |
| Q493R,N501Y | -2.93 | 0.92 | Q493R,N501Y |
| S494P,N501Y | -1.72 | 1.28 | S494P,N501Y |
| G496S | -0.23 | 0.20 | G496S |
| *N501Y | -1.68 | 1.21 | N501Y |

**Fig. 2.**
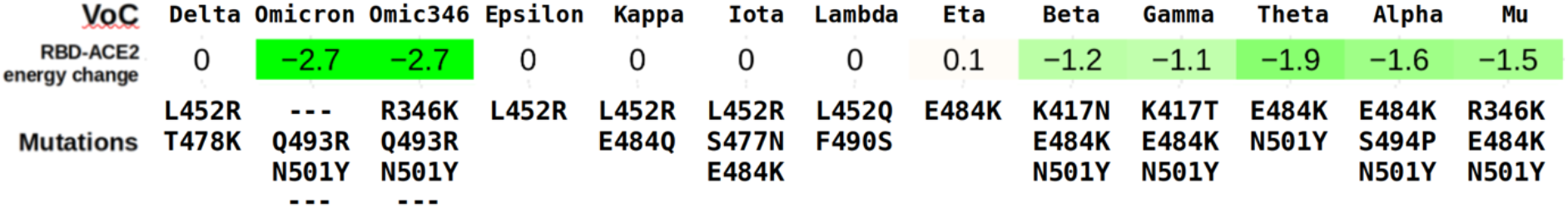
Illustration of predicted changes in binding energy (ddG) upon introduced mutations to the SARS-CoV-2 RBD-ACE-2 complexes, highlighting Omicron’s increased binding to ACE-2 (lower ddG). Numerical data for this illustration and other evaluated RBD mutants is provided in Table 1 below.

### Many publicly available neutralizing antibodies are predicted to have degraded binding affinity to the Omicron variant

To help address the question of whether Omicron has an increased ability to evade known SARS-CoV-2 neutralizing antibodies and possibly vaccine-induced protective response, we performed SFE predictions on 55 SARS-CoV-2 antibodies in complex with each of 13 selected SARS-CoV-2 VoC’s. Our predictions suggest that, in comparison with the Delta variant, Omicron can significantly degrade binding affinity for most of the analyzed Fabs (see first two columns in the Table 2 and Table 3 below). SFE-ddG results suggest that Omicron reduces affinity more than most of other variants. Negative values (colored in green) indicate improvements in binding in comparison to Wuhan SARS-CoV-2, and positive values (colored in orange) suggest weaker binding.

**Table 2.**
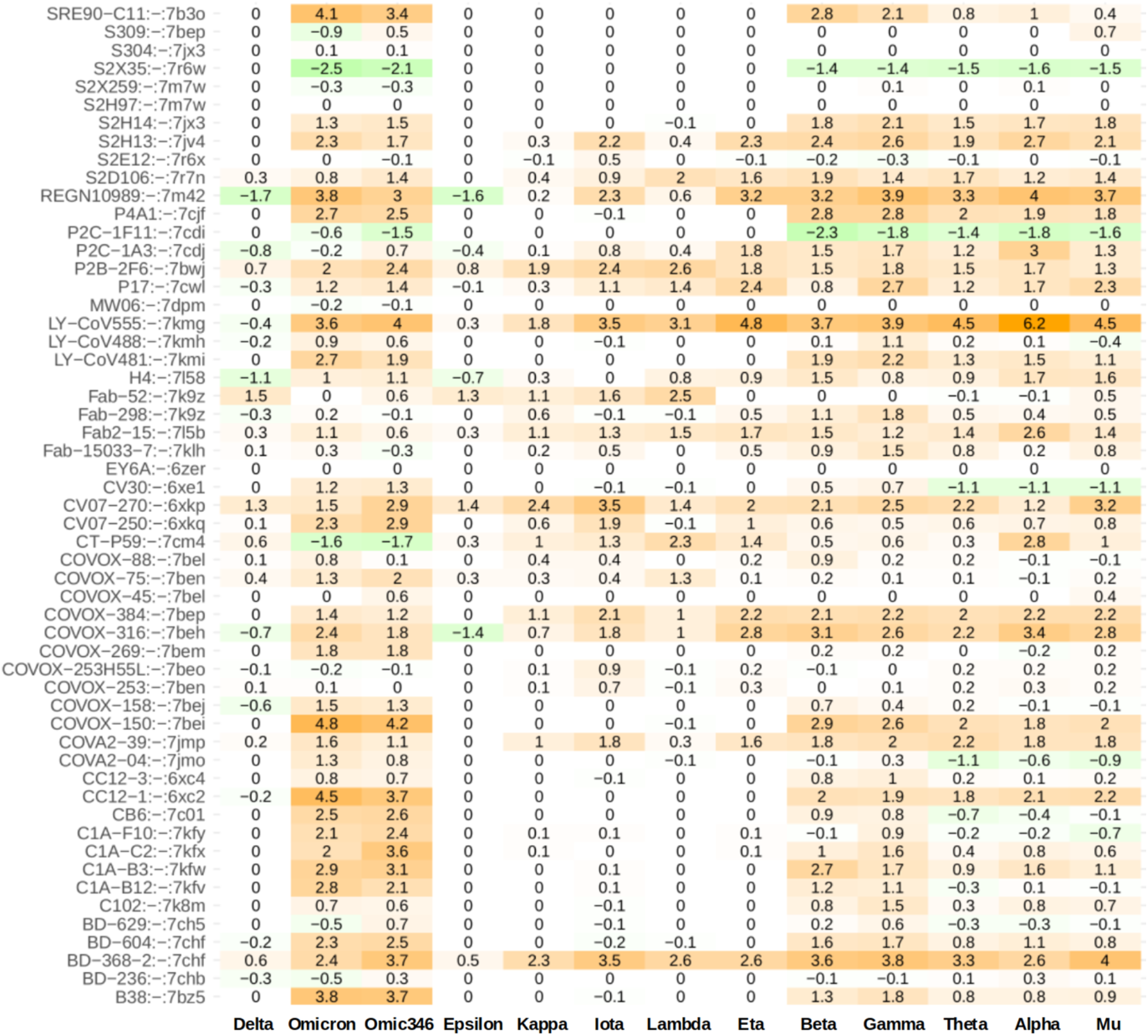
Predicted changes in energy (ddG estimates) upon introduction of mutations to SARS-CoV-2 RBD-Fab complexes. Fifty-five known neutralizing SARS-COV-2 monoclonal antibodies (Y-axis) were evaluated for ddG changes upon replacement of SARS-CoV-2 RBD with RBDs (X-axis) of 13 selected SARS-CoV-2 VoC.

**Table 3.** The impact on RBD-FAB binding of Omicron and Delta variants for 55 known SARS-CoV-2 neutralizing antibodies is compared by counting the number of estimated ddG scores that are above selected thresholds of 0.2, 0.5, and 1.0 (column-wise). Applying these ddG values as thresholds for viral escape, we observe that the Omicron variant may have an increased ability to evade binding with FABs over other variants, including Delta. In parentheses we provide the percentage of weakened antibodies from a total of 55 tested.

| **Variant** | **ddG>=0.2** | **ddG>=0.5** | **ddG>=1.0** |
| --- | --- | --- | --- |
| Delta: | 8 (15%) | 5 (9%) | 2 (4%) |
| Delta1: | 26 (47%) | 22 (40%) | 10 (18%) |
| Delta2: | 17 (31%) | 12 (22%) | 7 (13%) |
| Omicron: | 37 (67%) | 36 (65%) | 31 (56%) |
| Omic346: | 41 (75%) | 39 (71%) | 30 (55%) |

While positive predicted ddG values suggest a decrease in binding affinity, there is no clear threshold value at which binding is predicted to be abolished or significantly reduced. In Table 3 we present a comparison of Omicron versus other variants using three different such threshold values of ddGs. Results suggest that Omicron can significantly disrupt about 75% of neutralizing antibodies when a ddG threshold of 0.2 is considered as a breaking point (nearly twice as many as for Delta) or 56% of neutralizing antibodies when a threshold of 1.0 is used (more than three times as many as for Delta).

## Discussion

We structurally modeled the SARS-CoV-2 Omicron variant in-complex with ACE-2 and illustrated potentially significant changes in the RBD-ACE2 interface region. However, we caution that a reliable, experimentally-derived structure of the SARS-CoV-2 Omicron RBD is not currently available and the large number of mutations on Omicron introduces significant uncertainty of the model and our conclusions. Nevertheless, such analysis is particularly useful and informative while experimental structures the Omicron variant are forthcoming.

Our computational affinity predictions suggest the Omicron variant has increased affinity to ACE-2 compared to other Variants of Concern, which may have implications for increased infectivity, viral load, and/or transmissibility. We caution that affinity to ACE-2 is only one of many factors impacting transmissibility and infectivity, but these results suggest that experimental and epidemiological investigation is urgently needed. Previously published experimental results in [5] for some of the specific mutations present in Omicron are consistent with our predictions.

Our computational affinity predictions suggest that a large number of publicly known neutralizing antibodies will display significantly weaker binding to the Omicron variant compared to other VoC’s. We caution that binding strength is not fully predictive of the variant’s ability to evade vaccine-induced immunity and therapeutic antibodies. Moreover, our affinity predictions to ACE-2 and neutralizing antibodies are based on our predicted structure of Omicron and ACE-2, and *in silico* evaluation of binding interactions, which do not fully capture all aspects of the true real-world system. We also limited our analysis to single RBD-FAB or RBD-ACE-2 complexes (i.e., we did not evaluate the complete Spike protein nor trimer oligomeric states). Moreover, in the case of evaluated RBD-FAB complexes, we restricted analysis to only the FAB variable domains.

Despite these important caveats, we believe that computational predictions have an increasingly important role in rapid response to emerging, high consequence pathogens. Specifically, computational predictions can help quantitatively communicate risk and guide decision-making while awaiting experimental/real-world assessments. In ongoing and future work, we plan to compare our model predictions to experimental data to assess and improve our predictive capabilities.

## Acknowledgement

This work was supported by DARPA agreement number HR0011154580, Joint Program Execution Office agreement number 11647302, and Laboratory Directed Research and Development (LDRD 20-ERD-032) at Lawrence Livermore National Laboratory (LLNL). We would like to thank Christopher Earnhart, Amy Jenkins, Svetlana Hopkins, Tom Bates, and James Brase for helpful discussions and support.

This document was prepared as an account of work sponsored by an agency of the United States government. Neither the United States government nor Lawrence Livermore National Security, LLC, nor any of their employees makes any warranty, expressed or implied, or assumes any legal liability or responsibility for the accuracy, completeness, or usefulness of any information, apparatus, product, or process disclosed, or represents that its use would not infringe privately owned rights. Reference herein to any specific commercial product, process, or service by trade name, trademark, manufacturer, or otherwise does not necessarily constitute or imply its endorsement, recommendation, or favoring by the United States government or Lawrence Livermore National Security, LLC. The views and opinions of authors expressed herein do not necessarily state or reflect those of the United States government or Lawrence Livermore National Security, LLC, and shall not be used for advertising or product endorsement purposes.

Lawrence Livermore National Laboratory is operated by Lawrence Livermore National Security, LLC, for the U.S. Department of Energy, National Nuclear Security Administration under Contract DE-AC52-07NA27344.

